# Lysophosphatidic acid (LPA)-antibody (504B3) validation by free-solution assay and interferometry identifies off-target binding

**DOI:** 10.1101/2020.10.05.327122

**Authors:** Manisha Ray, Yasuyuki Kihara, Darryl J. Bornhop, Jerold Chun

## Abstract

Antibody specificity and sensitivity is required in basic and clinical research for ensuring scientific rigor and reproducibility, while off-target cross-reactivity could generate erroneous conclusions. Lysophosphatidic acid (LPA) is a bioactive lipid being targeted clinically by antibody strategies. Here, we reexamined binding properties of a commercially available monoclonal antibody (504B3) reported as specific for LPA using a free-solution assay measured in a compensated interferometric reader. The antibody showed comparable binding affinities to LPA and non-LPA lipids including phosphatidic acid (PA) and lysophosphatidylcholine (LPC). These results may alter conclusions drawn from current and past basic and clinical studies employing anti-LPA antibodies.

## Main Text

LPA is a potent, bioactive lipid that acts through six cognate G protein-coupled receptors (LPA_1-6_) [1, 2]. It is involved in many physiological processes such as cell proliferation, chemotaxis, and smooth muscle contraction, as well as cell survival [3]. Elevated LPA levels are associated with several disease pathologies, including cancer, hydrocephalus, and fibrosis [3-6], implicating therapeutic potential of modulating LPA pathways.

Lpath Inc. (merged with Apollo Endosurgery, Inc. in 2016) developed humanized monoclonal anti-LPA antibodies, including a phase 1a molecule, Lpathomab/LT3015 [7, 8]. The binding affinity and selectivity of Lpathomab/LT3015 was determined by enzyme-linked immunosorbent assay (ELISA) using unnatural, biotinylated lipid species [9], which showed nearly identical, nanomolar affinities to several LPA forms (14:0-, 18:0-, 18:1-, and 18:2-LPAs), without reported binding to other lipid species, including sphingosine 1-phosphate (S1P), 18:0-lysophosphatidylcholine (LPC), phosphatidic acid (PA), phosphatidylcholine (PC), and platelet-activating factor (PAF) [9]. A related anti-LPA antibody, 504B3, whose complementarity determining regions showed 79% identity to the Lpathomab/LT3015 [10], is commercially available (Echelon Biosciences, Product Number: Z-P200) and has been used for preclinical studies of spinal cord injury [11] and traumatic brain injury [12]. ELISA studies reported similar specificity and selectivity of 5043B compared to Lpathomab/LT3015 [11].

ELISA is a commonly used assay for determining binding affinity, but possesses a number of technical drawbacks that may impact accuracy. These include multiple washing steps that may under or overestimate affinities, immobilization that can increase non-specific interactions through conformational inflexibility [13, 14], and particularly use of biotinylated lipids that limit conformational flexibility of the lipids and introduce non-native epitopes [9]. By comparison, interferometric assays use unmodified, native binding partners such as native amino acids (serotonin, histamine, and dopamine) to their-specific antibodies [15], in free-solution and without need for labels on binding partners. FSAs use refractive index (RI) matched binding *sample* and *reference* solutions that when measured in a CIR, allow for determination of specific binding in real-time. The CIR detects the RI change that occurs as a result of binding-induced conformational and/or hydration changes produced by binding events in the *sample* compared to the null in the non-binding *reference*. The utility of FSA-CIR was recently demonstrated for high-sensitivity detection of native equilibrium binding K_D_s between LPAs and one of its receptors, LPA_1_ [16, 17].

FSA-CIR was used to reevaluate the equilibrium binding affinity (K_D_) of the anti-LPA mAb (504B3) against five phospholipids (18:1 LPA, 16:0 LPA, 18:1 LPC, 18:1 S1P, and 18:1-18:1 PA). The *sample* and *reference* were prepared by mixing ligand dilution series (0-100 nM; varied by ligand) with an equivalent volume of the 504B3 solution (*sample*; 10 μg/ml dissolved in PBS, pH 7.4) or PBS only (*reference*)(**Figure 1a**). The FSA signal (ΔRI; refractive index difference between *sample* and *reference*) as measured by the interferometer was plotted against ligand concentrations and showed nanomolar binding affinities of 504B3 to not only 18:1 LPA (K_D_ ≈ 3.73 ± 2.8 nM), but also to 18:1 LPC (K_D_ ≈ 8.5 ± 2.6 nM) and PA (K_D_ ≈ 3.3± 2.7 nM), with weaker affinity for 16:0 LPA **(Figure 1b; Table SI:1)**. Endogenous plasma concentrations of LPC (100-300 μm) range beyond 100X LPA concentrations (0.1-2 μm) [18-20], indicating that a vast majority of 504B3 and related antibodies used *in vivo* would be bound to LPC and PA rather than LPA. These results raise concerns about the use of this and related anti-LPA antibodies in basic and clinical research, such as in reports on protective effects in spinal cord injury [11] and traumatic brain injury [12]. More broadly, other antibodies generated against lipids and possibly other antigens may benefit from FSA-CIR over traditional methods like ELISA for the evaluation of molecular interactions under near-physiological conditions.

**Figure 1:**
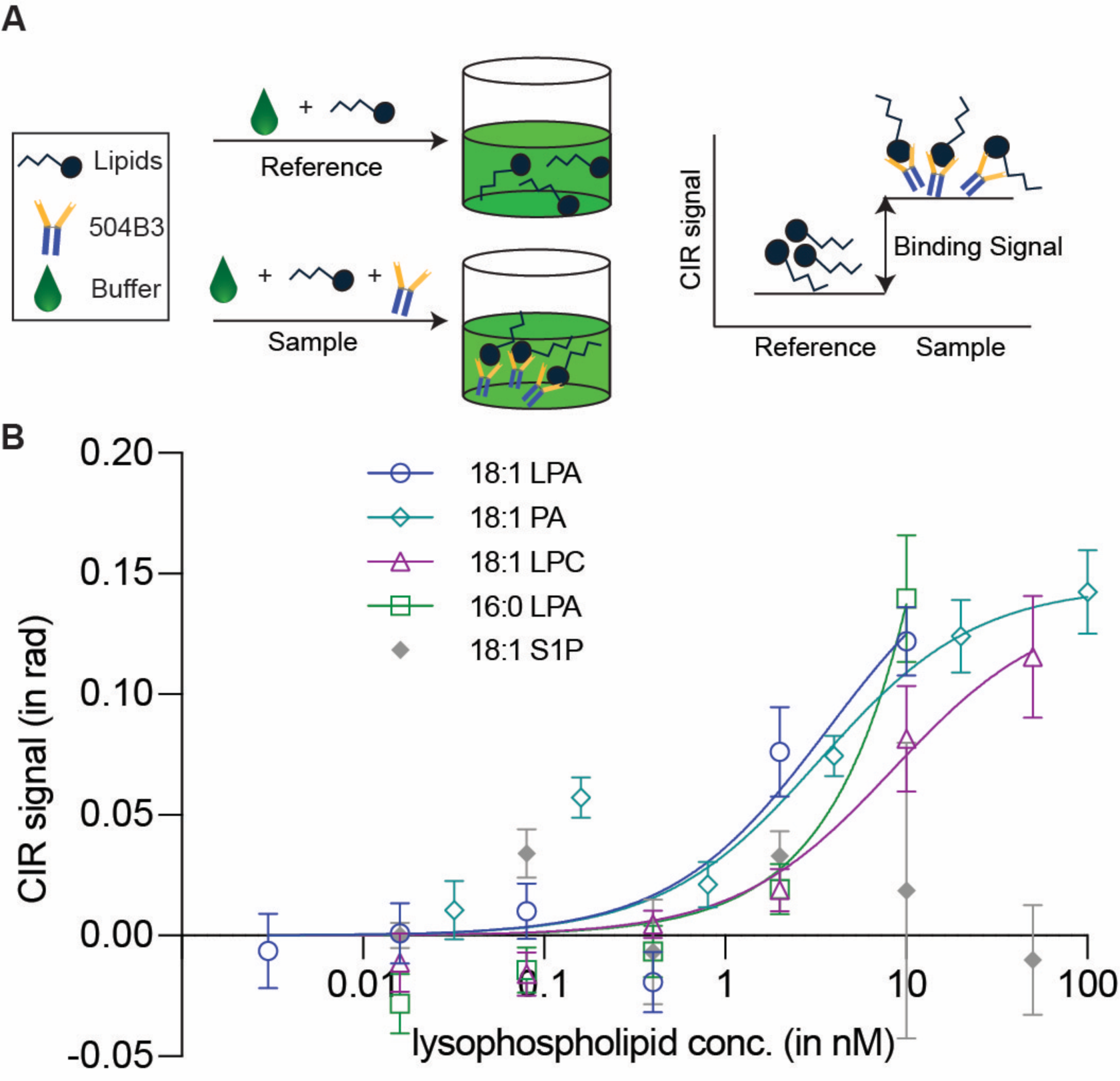
Free solution assay (FSA) measured by a compensated interferometric reader (CIR) to determine the binding constant (K_D_) for 504B3 anti-LPA antibody against five lysophospholipid forms. (a) FSA used to prepare RI matched sample and reference solutions. *See supplementary information for detailed preparation methods.* (b) Binding curves for each LP or PA ligand (*e.g.,* 18:1 LPA, 18:1 PA, 18:1 LPC, 16:0 LPA and 18:1 S1P) used to determine the binding constants (K_D_). Each graph shows an average of two independent binding isotherms (experimental replicates), each with quintuplicate measurements (technical replicates).

## Glossary

FSA: Free solution assay
CIR: Compensated Interferometric Reader
ELISA: Enzyme-linked immunosorbent Assay
LPA_1_: Lysophosphatidic Acid Receptor 1
LPA: Lysophosphatidic Acid
LPC: lysophosphatidylcholine
PA: Phosphatidic Acid
RI: Refractive Index

## Acknowledgements

The authors thank Mrs. Danielle Johns, Drs. Gwendolyn Kaeser and Laura Wolszon for their critical review and editing of this manuscript. Research funding was provided by the National Institutes of Health (R01NS084398 to JC) and the National Science Foundation (CHE-1610964 to DJB).

