## Supplementary material for "Lysophosphatidic acid (LPA)-antibody (504B3) validation by free-solution assay and interferometry identifies off-target binding": SI Files

**Supporting Information**

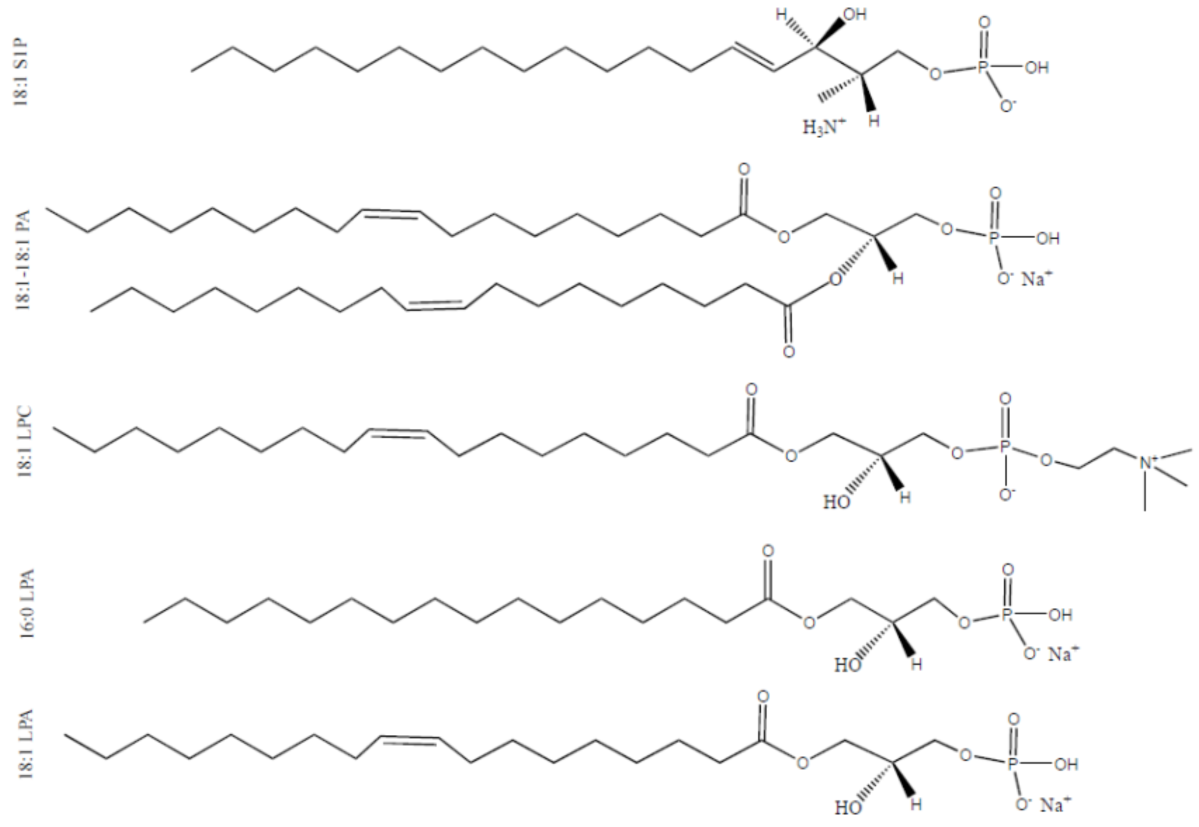


**Figure S1. Lysophospholipids (LPs) and phosphatidic acid (PA) used for binding assays with anti-LPA antibody 504B3 from Echelon Biosciences**. Lipids varied by acyl chain length (16:0, 18:1 LPA), head groups (18:1 S1P, 18:1 LPC) or the presence of a 2nd acyl chain (18:1-18:1 PA).

**Table S1. Binding constants ($K_D$) determined for different lysophospholipid (LP) forms and phosphatidic acid (PA) against the LPA antibody 504B3 from Echelon Bioscience**

| Lysophospholipid | $K_D$ from FSA ± SD | $B_{max}$, $R^2$ |
|---|---|---|
| 18:1 LPA | 3.7 ± 2.8 nM | 0.17; 0.83 |
| 18:1 PA | 3.3 ± 2.7 nM | 0.15; 0.81 |
| 18:1 LPC | 8.5 ± 2.6 nM | 0.14; 0.90 |
| 16:0 LPA | >30 nM | |
| 18:1 S1P | ~ 0 | |

**Materials and Methods:**

Lipids were purchased from Avanti polar lipids Inc. and the lysophosphatidic acid (LPA) antibody, 504B3, was purchased from Echelon biosciences.

**Lipid sample preparation**

Fresh lipid solutions were used in each binding assay. Powered lipids were reconstituted in solution: 18:1 LPA and 16:0 LPA were reconstituted in 1:1 (v/v) EtOH: $H_2O$; 18:1 LPC and 18:1-18:1PA were reconstituted in 50% EtOH/PBS; and 18:1 S1P was reconstituted in 0.4% BSA according to the protocol provided by Avanti Lipid Inc. for S1P.

**Free solution assay (FSA) configuration**

LPAs were delivered using fatty acid-free BSA for *in vivo* compatibility. A ligand dilution series (100, 50, 20, 4, 0.8, 0.16, 0.032, and 0 nM) was prepared in 0.01% BSA / 0.002% EtOH/PBS from an intermediate stock of 200 nM LPs in 0.01% BSA / 0.002% EtOH/PBS. Each LP dilution was combined with 1) PBS only to create the *reference* solution, or 2) 10 µg/ml of the 5043B antibody in PBS to create the binding *sample* solution. Solutions were kept at RT for approximately 1-hour to reach equilibrium and then analyzed using the compensated interferometric reader (**CIR**). The final concentrations were 5 µg/ml of antibody and 0-50 nM of ligand, in a final buffer composition of 0.005% BSA / 0.001% EtOH/PBS.

**CIR**

The details of the interferometer were described elsewhere (1, 2). Briefly, it consisted of a diode laser, two mirrors, one glass capillary and a CCD camera. A droplet train of *sample-reference* pairs was introduced and maintained at a constant flow rate in the capillary using a droplet generator (Mitos Dropix) and a syringe pump (flow rate 15µL/min). Coupling the interferometer with a syringe pump and a droplet generator resulted Compensated Interferometric Reader (CIR), a benchtop RI reader, where the FSA was measured (2, 3). The refractive index (RI) change between the binding *sample* and *reference* was measured as a positional shift in backscattered interference fringes produced from the interaction between an expanded beam profile of the laser and a capillary filled with sample-reference solutions (4). The shift of the backscattered fringe patterns is equivalent to molecular binding is quantified using fast Fourier transform of selected fringes captured in a CCD array. The assay was measured sequentially, as *reference*, then *sample*, and repeated seven times. The detailed procedure of the binding assay run in the interferometric reader was followed from our recently published $LPA_1$ receptor-LPA ligand binding assay (2).

1. Kammer MN, Kussrow AK, Olmsted IR, Bornhop DJ. A Highly Compensated Interferometer for Biochemical Analysis. ACS sensors. 2018;3(8):1546-52.
2. Ray M, Nagai K, Kihara Y, Kussrow A, Kammer MN, Frantz A, et al. Unlabeled lysophosphatidic acid receptor binding in free solution as determined by a compensated interferometric reader. Journal of lipid research. 2020;61(8):1244-51.
3. Kammer MN, Kussrow A, Gandhi I, Drabek R, Batchelor RH, Jackson GW, et al. Quantification of Opioids in Urine Using an Aptamer-Based Free-Solution Assay. Analytical chemistry. 2019;91(16):10582-8.
4. Kammer MN, Kussrow AK, Bornhop DJ. Longitudinal pixel averaging for improved compensation in backscattering interferometry. Optics letters. 2018;43(3):482-5.
